# Novel bir’s-foot trefoil RNA viruses provide insights into a clade of legume associated enamoviruses

**DOI:** 10.1101/368944

**Authors:** Humberto J. Debat, Nicolas Bejerman

## Abstract

Bird’s-foot trefoil (*Lotus corniculatus*) is a nutritious forage crop, employed for livestock foraging around the world. Here, we report the identification and characterization of two novel viruses associated with bird’s-foot trefoil. Virus sequences with affinity to enamoviruses (ssRNA (+); *Luteoviridae; Enamovirus*) and nucleorhabdoviruses (ssRNA (-); *Rhabdoviridae; Nucleorhabdovirus*) were detected in *L. corniculatus* transcriptome data. The tentatively named birds-foot trefoil associated virus 1 (BFTV-1) genome organization is characterized by 13,626 nt long negative-sense ssRNA. BFTV-1 presents in its antigenome orientation six predicted gene products in the canonical order 3′-N-P-P3-M-G-L-5′. The proposed birds-foot trefoil associated virus 2 (BFTV-2) 5,736 nt virus sequence presents a typical 5′-PO-P1-2-IGS-P3-P5-3′ enamovirus genome structure. Phylogenetic analysis suggests that BFTV-1 is closely related to *Datura yellow vein nucleorhabdovirus*, and that BFTV-2 clusters into a monophyletic cluster of legumes-associated enamoviruses. This sub-clade of highly related and co-divergent legume associated viruses provides insights on the evolutionary history of the enamoviruses. The bioinformatic reanalysis of SRA libraries deposited in the NCBI database constitutes an emerging approach to the discovery of novel plant viruses which should be important for both quarantine purposes and disease management.

---

From an agronomical point of view, bird’s-foot trefoil (*Lotus corniculatus* L.) is the most important *Lotus* species. This plant is a perennial legume mainly grown for fodder production in temperate regions and is considered one of the major forage legumes after lucerne (*Medicago sativa*) and white clover (*Trifolium repens*) [9]. It is a high quality forage that can be grazed or cut for hay or silage, and does not cause bloat in ruminants [13]. Intriguingly, to date the only virus which has been identified to affect this crop was the widely distributed alfalfa mosaic virus (AMV), in a field study of Prince Edward Island, Canada, more than 35 years ago [16].

Numerous novel viruses, many of them not inducing any apparent symptoms, have been identified from different environments using metagenomic approaches, which revealed our limited knowledge about the richness of a continuously expanding plant virosphere, widespread in every potential host assessed [18]. Genomic RNA molecules of these plant RNA viruses are often inadvertently co-isolated with host RNAs and their sequences can be detected in plant transcriptome datasets [15, 17]. In a recent consensus statement report, Simmonds et al [20] provide that viruses that are known only from metagenomic data can, should, and have been incorporated into the official classification scheme of the International Committee on Taxonomy of Viruses (ICTV). Thus, the analysis of public data constitutes an emerging source of novel *bona fide* plant viruses, which allows the reliable identification of new viruses in hosts with no previous record of virus infections. Here, we analyzed a transcriptomic dataset corresponding to bird’s-foot trefoil, available at the NCBI SRA database, which resulted in the identification and characterization of a novel enamovirus and nucleorhabdovirus associated with this crop.

The raw data analyzed in this study corresponds to an RNAseq NGS library (SRA: SRS271068), associated to NCBI Bioproject PRJNA77207. As described by Wang et al [22], it is derived of Illumina Hiseq 2000 sequencing of total RNA isolated from fresh flowers, pods, leaves, and roots from *L. corniculatus* collected in Qinling Mountain, Shaanxi Province, China (BioSample: SAMN00759026). The 26,492,952 2×90 nt raw reads from the SRA were pre-processed by trimming and filtering with the Trimmomatic tool as implemented in http://www.usadellab.org/cms/?page=trimmomatic, the resulting reads were assembled *de novo* with Trinity <u>v2.6.6 release</u> with standard parameters. *De novo* transcriptome assembly of SRA library SRS271068 [22] resulted in 55,473 transcripts, which were subjected to bulk local BLASTX searches (E-value < 1e-5) against a refseq virus protein database available at ftp://ftp.ncbi.nlm.nih.gov/refseq/release/viral/viral.1.protein.faa.gz. The resulting hits were explored by hand. A 6,524 nt transcript obtained a significant hit (E-value = 0, identity = 60%) with the L protein of datura yellow vein nucleorhabdovirus (DYVV; [6]). Further similarity analyses resulted in the detection of three more DYVV-like transcripts of 3,953 nt, 2,354 nt and 1,432 nt. The tentative virus contigs were curated by iterative mapping of reads using Bowtie2 http://bowtie-bio.sourceforge.net/bowtie2/index.shtml which was also employed for mean coverage estimation and reads per million (RPMs) calculations. The transcripts extended and polished by iterative mapping of raw reads were subsequently reassembled into a 13,626 nt long RNA contig with overall 57,7% sequence similarity with DYVV. This resulting RNA sequence was reconstructed with a total of 17,656 reads (mean coverage = 116.4X, reads per million (RPM) = 666.4. To further advance in the characterization of this sequence, virus annotation was instrumented as described in Debat [4], briefly, virus ORFs were predicted with ORFfinder (https://www.ncbi.nlm.nih.gov/orffinder/) domains presence and architecture of translated gene products was determined by InterPro (https://www.ebi.ac.uk/interpro/search/sequence-search) and the NCBI Conserved domain database v3.16 (https://www.ncbi.nlm.nih.gov/Structure/cdd/wrpsb.cgi). Further, HHPred and HHBlits as implemented in https://toolkit.tuebingen.mpg.de/#/tools/ was used to complement annotation of divergent predicted proteins by hidden Markov models.

The tentatively named birds-foot trefoil associated virus 1 (BFTV-1) genome organization is characterized by 13,626 nt long negative-sense single-stranded RNA (GenBank accession number MH614262) and contains six open reading frames (ORFs) in the anti-genome, positive-sense orientation (**Fig. 1A**). BLASTP searches identified these ORFs as potentially encoding: a nucleocapsid protein (N; ORF1, 462 aa), phosphoprotein (P; ORF2, 344 aa), movement protein (P3; ORF3, 325 aa), matrix protein (M; ORF 4, 294 aa), glycoprotein (G; ORF 5, 637 aa), and an RNA dependent RNA polymerase (L; ORF 6, 2104 aa), based on highest sequence identity scores with nucleorhabdoviruses, in particular with black currant-associated nucleorhabdovirus (BCaRV; [23]), DYVV and sonchus yellow net nucleorhabdovirus (SYNV; [12]) (**Table 1**). The coding sequences are flanked by 3’ leader (l) and 5’ trailer (t) sequences revealing a genome organization of 3’ l–N–P–P3–P4-M–G–L–t 5’ (**Fig.1A**), which is the basic organization described for plant rhabdoviruses [7] given that no accessory genes are encoded by the BFTV-1 genome. Like all plant rhabdoviruses, BFTV-1 genes are separated by intergenic “gene junctions” regions, which are composed of the polyadenylation signal of the preceding gene, a short intergenic region, and the transcriptional start of the following gene (**Supp. Table 1**). Interestingly, BFTV-1 consensus “gene junction” region sequence 3′-AUUCUUUUUGGUUGUA-5 ‘ is identical to that of BCaRV, DYVV and SYNV (**Supp. Table 1**). All BFTV-1-encoded proteins contain a classical mono or bi-partite nuclear localization signal (NLS) [7]. Importin-α dependent nuclear localization signals were predicted using cNLS Mapper available at http://nls-mapper.iab.keio.ac.ip/, the scores predicted an exclusively nuclear localization for N, M and L proteins, while for P, P3 and G the scores suggested localization to both the nucleus and cytoplasm (**Table 1**). Nuclear export signals were predicted using NetNES 1.1 available at www.cbs.dtu.dk/services/NetNES/ which predicted a leucine-rich nuclear export signal in BFTNRV N protein near amino acid position 350 (data not shown), suggesting both nuclear import and export. Similar results were reported for the predicted products of DYVV [7]. Amino acid sequence comparisons between the deduced BFTV-1 proteins and the corresponding sequences of other nucleorhabdoviruses revealed high similarity of BFTV-1 to BCaRV, DYVV and SYNV. BFTV-1 N, G and L proteins were the most identical to that encoded by BCaRV, DYVV and SYNV sharing a 37.0-58.4% similarity, whereas P, P3 and M were the more divergent (**Supp. Table 2**).

**Figure 1.**
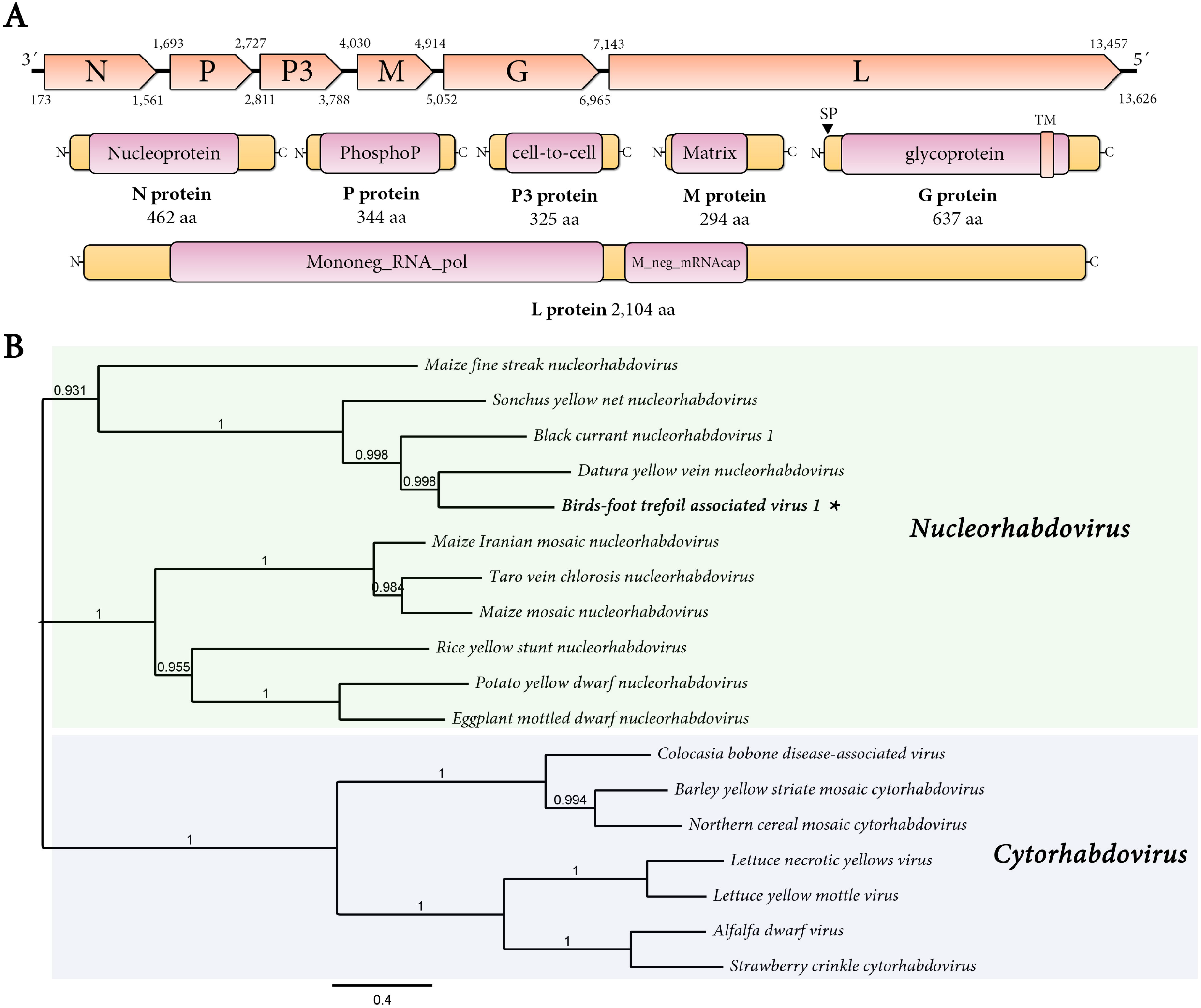
Structural characterization and phylogenetic insights of bird’s-foot trefoil associated virus 1 (BFTV-1) (**A**) Genome graphs depicting architecture and predicted gene products of BFTV-1. The predicted coding sequences are shown in orange arrow rectangles, start and end coordinates are indicated. Gene products are depicted in curved yellow rectangles and size in aa is indicated below. Predicted domains or HHPred best-hit regions are shown in curved pink rectangles. Abbreviations: N, nucleoprotein CDS; P, phosphoprotein CDS; P3, putative cell-to-cell movement protein CDS; M, matrix protein CDS; G, glycoprotein CDS; L, RNA dependent RNA polymerase CDS; TM, trans-membrane domain; SP, signal peptide. (**B**) Maximum likelihood phylogenetic tree based on amino acid alignments of the L polymerase of BFTNRV and other plant rhabdoviruses. The tree is rooted at the midpoint; nucleorhabdovirus and cytorhabdovirus clades are indicated by green and blue rectangles, respectively. The scale bar indicates the number of substitutions per site. Node labels indicate FastTree support values. The viruses used to construct the tree, and their accession numbers are as follows: black currant nucleorhabdovirus (MF543022), alfalfa dwarf virus (KP205452), barley yellow striate mosaic cytorhabdovirus (KM213865), colocasia bobone disease associated-virus (KT381973), datura yellow vein nucleorhabdovirus (KM823531), eggplant mottled dwarf nucleorhabdovirus (NC_025389), lettuce yellow mottle virus (EF687738), lettuce necrotic yellows virus (NC_007642); maize fine streak nucleorhabdovirus (AY618417), maize Iranian mosaic nucleorhabdovirus (DQ186554), maize mosaic nucleorhabdovirus (AY618418), northern cereal mosaic cytorhabdovirus (AB030277), potato yellow dwarf nucleorhabdovirus (GU734660), rice yellow stunt nucleorhabdovirus (NC_003746); sonchus yellow net virus (L32603), taro vein chlorosis virus (AY674964) and strawberry crinkle cytorhabdovirus (MH129615).

Phylogenetic insights based on predicted replicase proteins were generated by MAFTT 7 https://mafft.cbrc.jp/alignment/software/ multiple amino acid alignments (BLOSUM62 scoring matrix) using as best-fit algorithm E-INS-i (BFTV-1) or G-INS-I (BFTV-2). The aligned proteins were subsequently used as input for FastTree 2.1.5 (http://www.microbesonline.org/fasttree/) maximum likelihood phylogenetic trees (best-fit model = JTT-Jones-Taylor-Thorton with single rate of evolution for each site = CAT) computing local support values with the Shimodaira-Hasegawa test (SH) and 1,000 tree resamples. The obtained tree shows that BFTV-1 clusters together with other viruses within the genus *Nucleorhabdovirus* (**Fig.1B**). BFTV-1 appears to have a close evolutionary relationship with BCaRV, DYVV and SYNV, which share a similar genomic organization to that described for BFTV-1 [8, 13, 23]. It is tempting to suggest that BFTV-1, BCaRV, DYVV and SYNV may represent the most ancestral clade within nucleorhabdoviruses, since no accessory genes are present in their genomes. Taken together our results suggest that BFTV-1 should be considered a member of a new virus species in the genus *Nucleorhabdovirus.*

In addition to the nucleorhabdovirus, the BLASTX searches of the obtained transcriptome showed a 5,622 nt transcript which obtained a significant hit (E-value = 0, identity = 74%) with the P1-P2 fusion protein of Alfalfa enamovirus 1 (AEV-1; [2]). Mapping of the SRA raw reads extended the contig into a 5.736 nt RNA sequence (mean coverage = 61.6X, reads per million (RPM) = 147.1) with an overall identity of 69.2% with AEV-1. The assembled sequence of bird’s-foot trefoil virus 2 (BFTV-2) consists of 5.736 nt (GenBank accession number MH614261) and presents a typical 5′-PO-P1-2-IGS-P3-P5-3′ enamovirus genome structure (**Fig.2A**).

**Figure 2.**
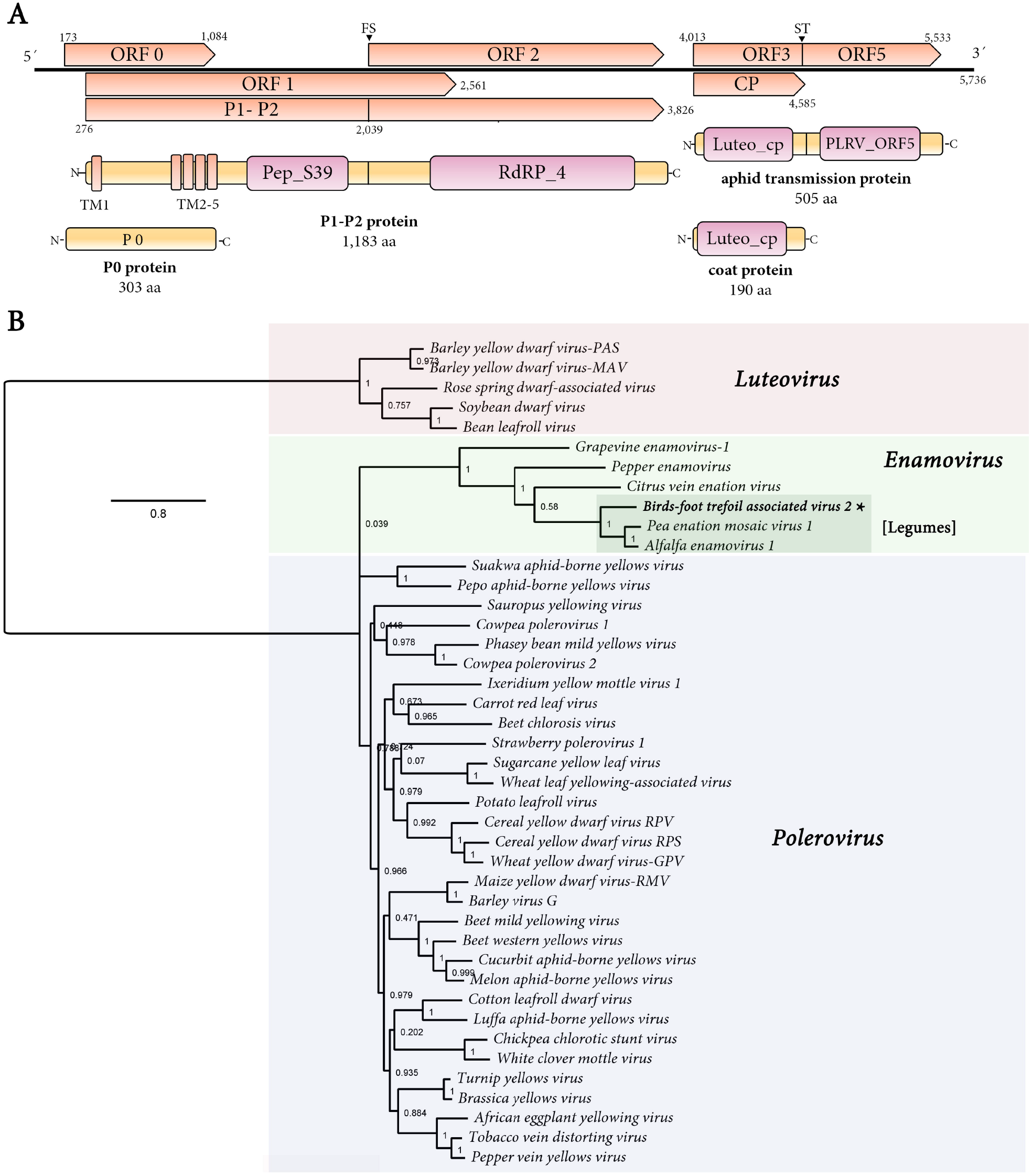
Structural characterization and phylogenetic insights of birds-foot trefoil virus 2 (BFTV-2) (**A**) Genome graphs depicting architecture and predicted gene products of BFTV-2. The predicted coding sequences are shown in orange arrow rectangles, start and end coordinates are indicated. Gene products are depicted in curved yellow rectangles and size in aa is indicated below. Predicted domains or HHPred best-hit regions are shown in curved pink rectangles. Abbreviations: CP, coat protein CDS; P1-P2, RNA dependent RNA polymerase fusion protein CDS; FS, −1 ribosomal frameshifting signal; TM1-5, transmembrane domains; ST, signal of translation read-through of a UGA stop codon; Pep_S39, peptidase S39 domain; RdRP_4, RdRP domain; Luteo_cp luteovirid coat protein domain; PLRV_ORF5, polerovirus readthrough protein domain. (**B**) Maximum likelihood phylogenetic tree based on amino acid alignments of the P1-P2 polymerase of BFTV-2 and other enamoviruses. The *Luteovirus* genus was used as tree root; *Enamovirus, Polerovirus* and *Luteovirus* clades are indicated by green, blue, and pink rectangles, respectively. The scale bar indicates the number of substitutions per site. Node labels indicate FastTree support values. The viruses used to construct the tree and their accession numbers are provided as **Supplementary Table 3**.

The first ORF, ORF 0, consists of 909 nt encoding a putative P0 of 303 aa with a calculated molecular mass of 33.8 kDa. In enamoviruses, P0 has been shown to function as an RNA silencing suppressor [11]. BFTV-2 P0 is more similar to that of legume-infecting enamoviruses AEV-1 and pea enation mosaic virus-1 (PEMV-1), where an F-Box-like motif located on the N-region of the predicted protein was identified (**Supp. Fig. 1A**). Interestingly, this motif was not identified in the non-legume infecting enamoviruses. The F-box-like motif is involved in silencing suppression [11], therefore it is tempting to speculate that other encoded proteins, rather than P0, may be involved in silencing suppression of the non-legume infecting enamoviruses. Nevertheless, proper assays should be employed to test this hypothesis. The second ORF, ORF1, which contains 2,283 nt, is predicted to be expressed by a ribosomal leaky scanning mechanism for a protein P1 (761 aa, 83.9 kDa). The third ORF, ORF2, which is translated by a −1 ribosomal frameshift from ORF 1 overlaps ORF1 at its 5’ end and is predicted to produce an ORF1-ORF2 fusion protein, P1-P2 (1183 aa, 131.4 kDa). The canonical motif for a −1 frameshift site is X_XXY_YYZ. A putative slippery sequence was detected at position 2,039 of the type G_GGA_AAC (**Supp. Fig. 2A**), identical to that of PEMV-1 [5]. In addition, an H-type highly structured (Free energy = −12.80) 40 nt pseudoknot was detected eight nt downstream (seven nt spacer) immediately following the slippery sequence (**Supp. Fig. 2B**) which was predicted with the KnotInFrame tool available at https://bibiserv2.cebitec.uni-bielefeld.de/knotinframe and visualized with the VARNA 3.93 applet (http://varna.lri.fr/). It is worth noting that the frameshifting pseudoknot in other polero/enamoviruses, including PEMV-1, is seven nt from the end of the shifty site (6 nt spacer), given that the spacing between the slippery sequence and pseudoknot is determinant [1] further studies should confirm the tentative prediction of this report. P1 and P1-P2 have a putative involvement in virus replication, while P1 is considered a serine-like protease, and the frameshift region (P2) of the P1-P2 protein is thought to contain the core domains of the viral RNA-dependent RNA polymerase (RdRP) [5]. A serine-protease-like domain (peptidase S39, pfam02122, P1 residues 314-515; E-value, 3.75e-42) in P1, and an RdRP domain (RdRP-4, pfam02123) resembling those from members of the genera *Polerovirus, Rotavirus* and *Totivirus* in P2 (P1-P2 residues 734-1116; E-value, 2.26e-61) were found when these protein sequences were analyzed (**Fig. 2A**). The fourth ORF, ORF3, consists of 570 nt, encoding a predicted coat protein P3 (190 aa, 21.2 kDa). The fifth ORF, ORF5, is a putative in-frame read-through product of ORF3 encoding a fusion protein P3-P5 (506 aa, 56.1 kDa). BFTV-2 presents the canonical stop-codon context for read-through of enamovirus (UGA-GGG; [9]). Moreover, between the 4,601-6,665 nt coordinates, 15 nt downstream the stop codon, 11 CCNNNN tandem repeat motif units were found, which are associated with ORF 3 stop codon read-through [3]. Furthermore, signatures in a cytidine-rich domain immediately after the stop codon (5’C-rich) and a branched stem-loop structure 684 nt downstream of the CP stop codon (CD-DRTE), which are required for ORF3 readthrough [24], were also identified in the BFTV-2 genomic sequence (**Supp. Fig. 3A-B**). Both BFTV-2 5’C-rich and CD-DRTE regions showed high identity with that of AEV-1, with the expected complementary bases of the two regions identical with the exception of a G-to-A SNP at the CD-DRTE, which is a non-complementary base in both BFTV-2 and AEV-1. P3 is the coat protein (CP), whereas the CP read-through extension (P5) of P3-P5 is thought to be an aphid transmission subunit of the virus [5]. While the CP region of the predicted P3-P5 protein is more conserved in enamovirus, some emerging motifs could be observed, such as a proline rich stretch immediately following the read-through region (**Supp. Fig. 1.B**). It is tempting to speculate that this motif could be involved in the transmission of BFTV-2; nonetheless, mutagenesis analysis coupled with virus transmission assays should be conducted to test this hypothesis. When these protein sequences were analyzed in detail, a luteovirid CP domain (Luteo_coat, pfam00894) in P3 (P3 residues 55-188; E-value, 1.63e-70) and a polerovirus read-through protein domain (PLRV_ORF5, pfam01690) in P5 (P3-P5 residues 221-424; E-value, 1.16e-48) were identified.

The functional and structural results indicate that BFTV-2 represents the genome sequence of a virus that should be taxonomically classified in the family *Luteoviridae.* Viruses in the family *Luteoviridae* contain a single-stranded positive-sense RNA genome and are classified into three genera, *Enamovirus, Luteovirus* and *Polerovirus.* Unlike poleroviruses, enamoviruses do not encode a P4 movement protein, and luteoviruses lack a P0 gene [8]. Therefore, BFTV-2 appears to be an enamovirus because its genomic structure, characterized by presenting a P0 gene, and not encoding a P4 protein. The BFTV-2 ORFs were compared to the predicted ORFs of AEV-1, PEMV-1, grapevine enamovirus-1 (GEV-1; [19]) and those of citrus vein enation virus (CVEV; [21]), in order to determine nucleotide and deduced amino acid sequence identities (**Table 1**). The maximum nt sequence identity for any gene CDS was 75.4, 77.1, 47.2 and 44.9%, respectively, whereas the maximum aa sequence identity for any gene product was 82.6, 85.3, 44.0 and 37.7%, respectively (**Table 1**). Therefore, the differences in aa sequence identity for each gene product were greater than 10%, which is one of the criteria used by the ICTV to demarcate species in the genera *Polerovirus* and *Luteovirus* [8] and also in *Enamovirus.* Consequently, BFTV-2 may belong to a new species in the genus *Enamovirus.* Further, in a phylogenetic tree based on the P1-P2 fusion protein aa sequence of viruses of the family *Luteoviridae*, BFTV-2 clustered with AEV-1 and PEMV-1 in a monophyletic clade of legumes-associated enamoviruses, within the enamovirus complex (**Fig. 2B**), which suggest that all legume-infecting enamoviruses share a common ancestor. Further assessment of the genetic distance of each gene product of all reported enamoviruses (**Supp. Fig. 4**) indicates an evident high similarity among the legume associated enamoviruses, including BFTV-2. It is interesting that BFTV-2 in phylogenetic trees (**Fig. 2B**) clusters more closely with AEV-1 than PEMV-1. Unlike PEMV-1 which was first reported in pea, AEV-1 has been detected only in alfalfa, a member of the *Faboideae* sub-family of legumes, shared with bird’s foot trefoil. Thus, these legume viruses appear to consistently co-diverge among them. Additional legume viruses would be helpful to entertain whether the evolutionary path of enamoviruses is characterized by co-divergence. AEV-1 is associated to a complex of viruses responsible of the alfalfa dwarf disease [2]. It is interesting that BFTV-2 was detected in concert with a nucleorhabdoviruses. Future studies should focus on determining if there might be any synergistic effect on the host caused by the co-infection of enamoviruses and nuclerhabdoviruses: a subtle interaction scenario that may have important implications in management of associated diseases. For instance, the umbravirus PEMV-2 provides the movement protein [5, 8] function lacking in the enamovirus PEMV-1; as no umbraviruses were identified in the bird’s-foot trefoil dataset, therefore it is tempting to speculate that perhaps the identified nucleorhabdovirus could be providing the movement functions to complement and thus allow in concert the movement and replication of BFTV-2.

In summary, the analysis of public SRA data constitutes an emerging source of novel plant viruses. Using this approach, we report the identification and molecular characterization of two novel viruses associated with bird’s-foot trefoil. Our analyses unveiled that these viruses may be considered novel species of the genera *Nucleorhabdovirus* (BFTV-1) and *Enamovirus* (BFTV-2). The RNA viruses reported here provide for the first time a glimpse of the virus landscape of this important crop. More importantly, these novel viruses provide evidence of a sub-clade of highly related and co-divergent legume associated enamoviruses as new clues on the understanding of the evolutionary history of this genus of viruses. Future studies should assess the prevalence of BFTV1 and BFTV-2, and unravel whether the infection of these novel viruses is associated to specific symptoms.

## Acknowledgments

We would like to express a sincere gratitude to the generators of the underlying data used for this work: Dr. Ying Wang and Dr. Zhezhi Wang. By following open access practices and supporting accessible raw sequence data in public repositories available to the research community, they have promoted the generation of new knowledge and ideas.

## Compliance with ethical standards

### Ethical Approval

This article does not contain any studies with human participants or animals performed by any of the authors.

### Funding

This research did not receive any specific grant from funding agencies in the public, commercial, or not-for-profit sectors.

---

**Supplementary Figure 1.**
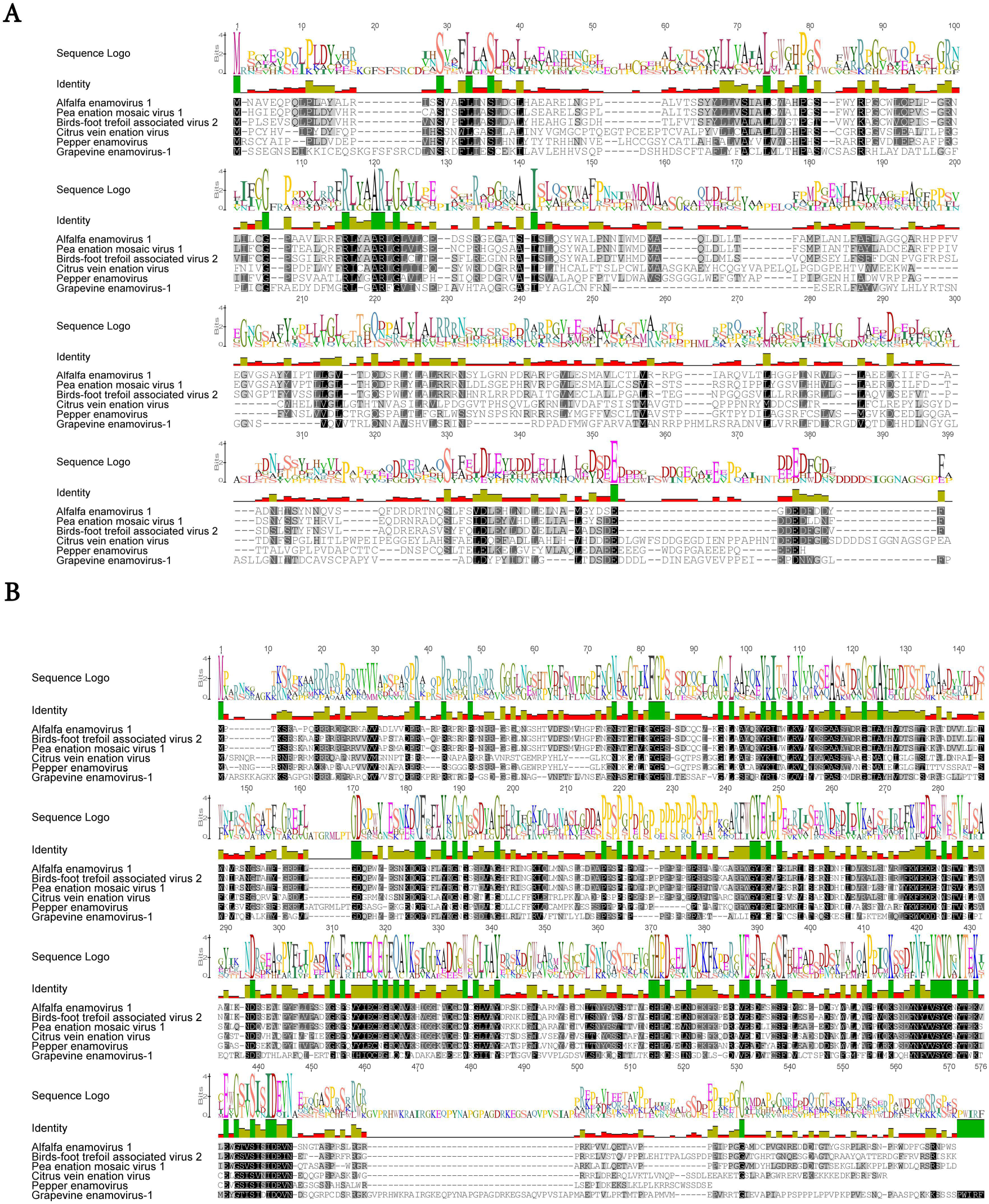
Amino acid sequence alignments of P0 protein of enamoviruses (**A**) and P3-P5 readtrough aphid transmission predicted protein of enamoviruses (**B**). Sequence similarity is depicted form white to black and sequence logo expressed as bits per position.

**Supplementary Figure 2.**
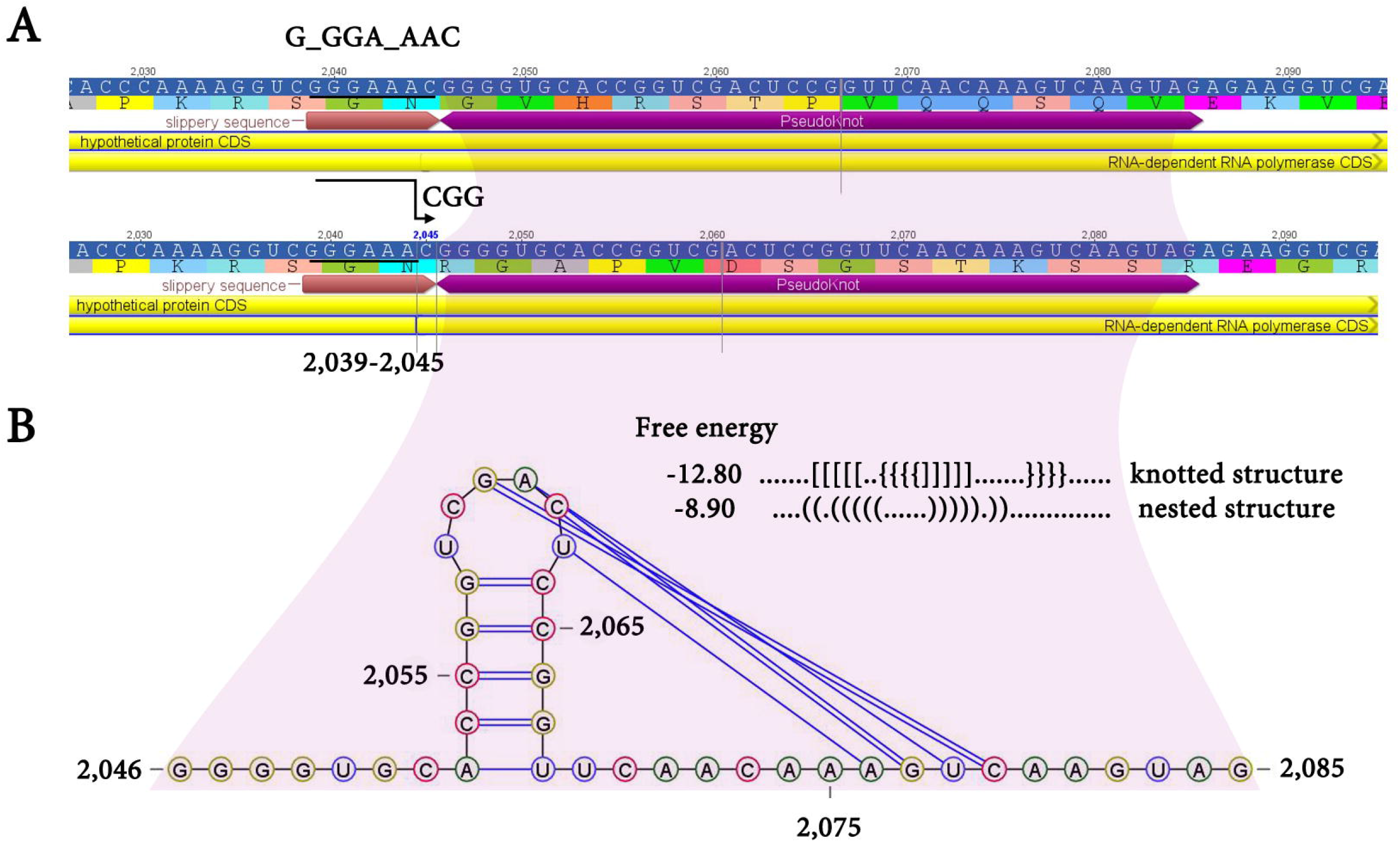
Ribosomal frameshifting prediction of P1-P2 CDS region. (**A**) Canonical motif for a −1 frameshift site of the form X_XXY_YYZ. The putative slippery sequence was detected at position 2,039 of the type G_GGA_AAC, identical to that of pea enation mosaic virus-1 (PEMV-1). (**B**) H-type highly structured (Free energy = −12.80) 40 nt pseudoknot detected starting 8 nt downstream the slippery sequence.

**Supplementary Figure 3.**
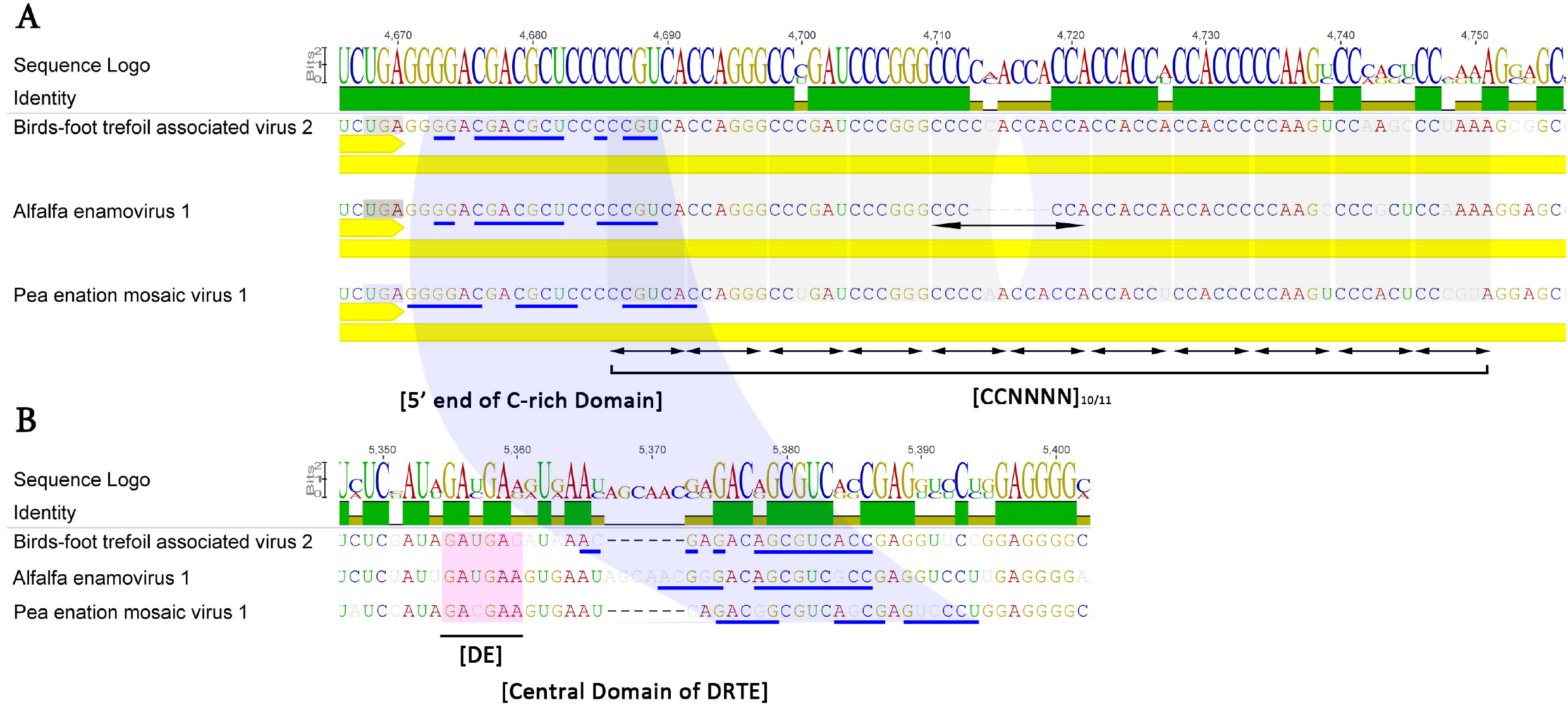
CCNNNN oligomer repeats and long-range tentative base pairing between the 5’ end of C-rich domain adjacent to the leaky stop codon of ORF3 (**A**) and the distal readthrough element (DRTE) of BFTV-2 and other legume associated enamoviruses (**B**). CCNNNNs are indicated by arrows, the distal element motif (DE) is depicted by a pink rectangle, complementary bases are underlined in blue.

**Supplementary Figure 4.**
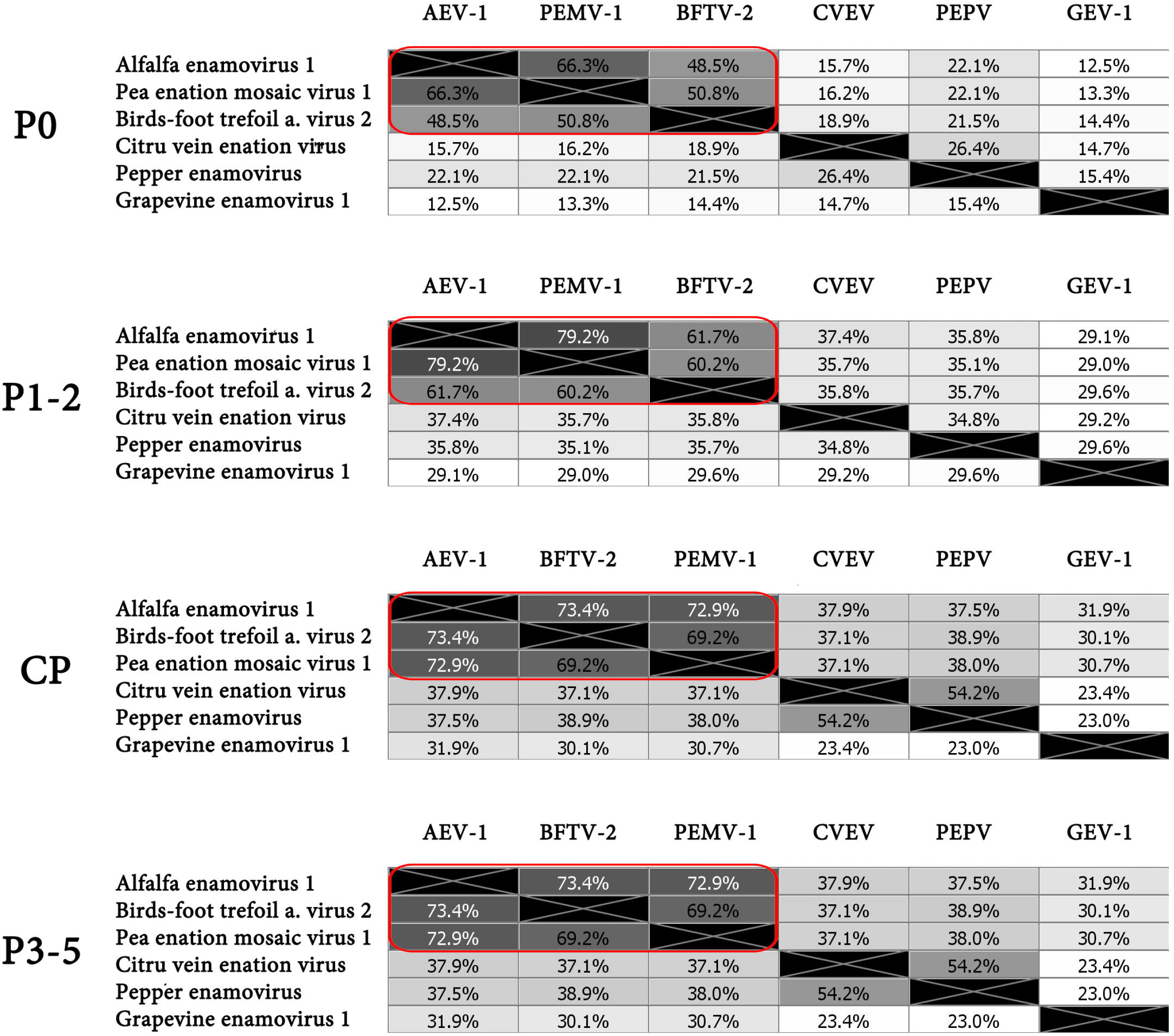
Genetic distances (% aa identity) of predicted gene products of BFTV-2 and reported enamoviruses support a sub-clade of legume associated enamoviruses (circled in red). Values are also presented as colored heatmaps ranging from 0% (white) to 100% (black) identity. AEV-1, alfalfa enamovirus 1; PEMV-1, pea enation mosaic virus-1; CVEV, citrus vein enation virus; PEPV, pepper enamovirus; GEV-1, grapevine enamovirus-1.

